# Chemical inhibition of the integrated stress response impairs the ubiquitin-proteasome system

**DOI:** 10.1101/2024.03.13.584747

**Authors:** Shanshan Xu, Maria E. Gierisch, Enrica Barchi, Ina Poser, Simon Alberti, Florian A. Salomons, Nico P. Dantuma

## Abstract

The Integrated Stress Response Inhibitor (ISRIB) is an experimental compound that has been used to explore the potential beneficial effects of reducing the activation of the integrated stress response (ISR). As the ISR is a protective response, there is, however, a risk that its inhibition may compromise the cell’s ability to restore protein homeostasis. Here, we show that ISRIB treatment impairs degradation of proteins by the ubiquitin-proteasome system (UPS) during proteotoxic stress in the cytosolic, but not nuclear, compartment. Degradation of proteins intercepted by ribosome quality control (RQC) was particularly affected as accumulation of a UPS reporter substrate for ribosome quality control (RQC) was comparable to the level observed after proteasome inhibition. Consistent with impaired RQC, ISRIB treatment caused an accumulation of polyubiquitylated and detergent insoluble defective ribosome products (DRiPs) in the presence of puromycin. As depletion of the RQC ubiquitin ligase listerin partially restored ubiquitin-dependent proteasomal degradation, these data suggest that the persistent protein translation during proteotoxic stress in ISRIB-treated cells increases the pool of newly synthesized proteins targeted by RQC, which aggravates UPS dysfunction by overloading the cytosolic UPS.

## Introduction

The integrated stress response (ISR) is a conserved signaling network that couples cellular stress detection to protein synthesis with as final goal to maintain protein homeostasis^1^. Extracellular and intracellular stress conditions can activate the ISR through stress-sensing kinases that phosphorylate serine residue 51 of the eukaryotic initiation factor 2α (eIF2α) subunit^2,3^. Eukaryotic initiation factor 2 (eIF2) is a heterotrimeric GTPase composed of α, β, and γ subunits, which form a ternary complex with GTP and the methionine initiator tRNA to initiate protein translation^4^. Once the start codon is decoded, GTP is hydrolyzed to GDP by the guanine nucleotide exchanging factor eIF2B, and eIF2 is released from the ribosome, forming a new ternary complex to engage in another round of translation initiation^5^. Phosphorylated eIF2α acts as a competitive inhibitor of eIF2B nucleotide exchanging activity, thereby limiting the recycling of eIF2, which results in reduced translation initiation and protein synthesis^6^. The untranslated mRNA in complex with disassembled 40S ribosome subunits interacts with RNA binding proteins and undergoes phase separation giving rise to cytosolic stress granules, which are membrane-less structures that function as temporary storage deposits for untranslated mRNAs^7^.

While the ISR is essentially cytoprotective and increases cell viability and survival during challenging conditions, its activation has also been linked to various pathological conditions, including cancer, diabetes, and neurodegenerative diseases^1^. The prevalence of ISR activation in these diseases has motivated the search for drugs that can inhibit or block this signaling cascade and prevent possible damage caused by the aberrant induction of this response. The Integrated Stress Response Inhibitor (ISRIB) is an experimental compound originally identified in a high-throughput screen for inhibitors that selectively mitigate PERK signaling^8^. Further studies revealed that ISRIB inhibits the ISR by allosterically destabilizing the complex of eIF2B with phosphorylated eIF2α^9^. By preventing the inhibitory effect of phosphorylated eIF2α on eIF2B, ISRIB blunts the ISR’s ability to block protein synthesis and to induce the formation of stress granules^10^. Thus, in the presence of ISRIB, protein synthesis persists while stress granules are not formed in response to proteotoxic stress.

It has been proposed that pharmacological inhibition of the ISR may be beneficial in the context of neurodegenerative diseases that are linked to activation of this pathway^11^. Indeed, ISRIB has been shown to enhance cognitive memory in prion-diseased mice, arguing for therapeutic applications of ISRIB in a clinical setting^12^. Considering the protective nature of the ISR, adverse effects may occur, but there is currently little insight into the negative consequences of curtailing this response. Induction of ER stress sensitizes cancer cells to ISRIB toxicity, consistent with a compromised ability of cells to deal with stress conditions^8,13^. This synergistic effect may allow selective targeting of cancer cells^14^, but it may also undermine its applicability in neurodegenerative disease as it could exacerbate cellular pathologies that involve ER stress^15^. Moreover, in other studies it has been reported that prolonging the ISR has a therapeutic effect in mouse models for protein-misfolding diseases^16,17^. While these findings are not mutually exclusive, it raises questions on how inhibition of the ISR affects other protein quality mechanisms in the cell.

The ubiquitin-proteasome system (UPS) is the primary pathway for the elimination of misfolded proteins during proteotoxic stress^18,19^. In essence it consists of a two-step process in which misfolded proteins are first modified by the small protein modifier ubiquitin and next degraded by the proteasome complex, which selectively hydrolyses ubiquitylated proteins^20^. While the ISR aims at restoring proteostasis by preventing the synthesis of misfolded proteins, the UPS contributes to this effort through the elimination of already existing misfolded proteins through proteolytic destruction. Thus, these two processes join efforts in restoring and maintaining protein homeostasis by reducing protein synthesis and stimulating degradation^21^. This raises the question as to whether the persistent protein synthesis in ISRIB-treated cells challenges the UPS during proteotoxic stress by increasing the load of misfolded substrates.

We report that ISRIB treatment negatively affects the UPS and compromises its ability to clear ubiquitylated substrates in cells undergoing proteotoxic stress. Consistent with a causal link with protein synthesis, ISRIB affects primarily the UPS in the cytosolic compartment and causes an increase in polyubiquitylated and insoluble defective newly synthesized proteins (DRiPs). Our data suggest that the persistent synthesis of proteins under conditions that compromise protein folding impairs the UPS and may thereby generate an intracellular environment that favors protein aggregation, a condition intrinsically connected to age-related neurodegenerative diseases.

## Results

### ISRIB aggravates UPS impairment during proteotoxic stress

We have previously shown that induction of proteotoxic stress by thermal stress transiently impairs the functionality of the UPS, resulting in the accumulation of polyubiquitylated substrates^22^. In this study, we used a mild heat shock as an experimental paradigm for an instant, reversible proteotoxic stress that inhibits protein synthesis and induces the formation of cytosolic stress granules^23^. To simultaneously access the formation of stress granules and the functional status of the UPS, we generated a human melanoma MelJuSo cell line that stably expresses mCherry-tagged G3BP1, a marker for stress granules^24^ and the ubiquitin fusion degradation (UFD) substrate ubiquitin^G76V^-yellow fluorescent protein (Ub-YFP), a sensor for ubiquitin-dependent proteasomal degradation^25^. Exposure of the reporter cell lines to a mild heat shock induced the formation of cytosolic stress granules as evidenced by the punctate cytosolic localization of mCherry-G3BP1 (**Fig. 1a, upper panels**). Importantly, induction of stress granule was efficiently blocked by ISRIB treatment (**Fig. 1a, upper panels**), which is in line with the reported biological activity of ISRIB on stress granule formation^8^. Exposure to heat shock induced phosphorylation of eIF2α in the absence and presence of ISRIB (**Fig. S1**), consistent with induction of the ISR by thermal stress and the fact that ISRIB inhibits the ISR downstream of eIF2α phosphorylation^9^. The Ub-YFP reporter substrate accumulated in the aftermath of the proteotoxic insult, indicating inefficient clearance of ubiquitylated proteins by proteasomal degradation in untreated and ISRIB-treated, thermally stressed cells (**Fig. 1a, lower panels**). Accumulation of Ub-YFP in response to heat shock is consistent with our earlier finding that mild thermal stress has a negative impact on ubiquitin-dependent proteasomal degradation^22^. Notably, quantitative analysis of the Ub-YFP levels 4 hours after heat shock showed that ISRIB treatment caused a significant increase in the accumulation of the UFD reporter as compared to untreated, thermally stressed cells (**Fig. 1b**). The increase in ISRIB-treated cells corresponded to approximately 10% of the increase in the Ub-YFP level that was obtained when the UPS was fully blocked with 200 nM of the proteasome inhibitor epoxomicin (**Fig. S2**). A time-course analysis confirmed that the Ub-YFP accumulated to higher levels in ISRIB-treated as compared to untreated cells (**Fig. 1c**).

**Figure 1.**
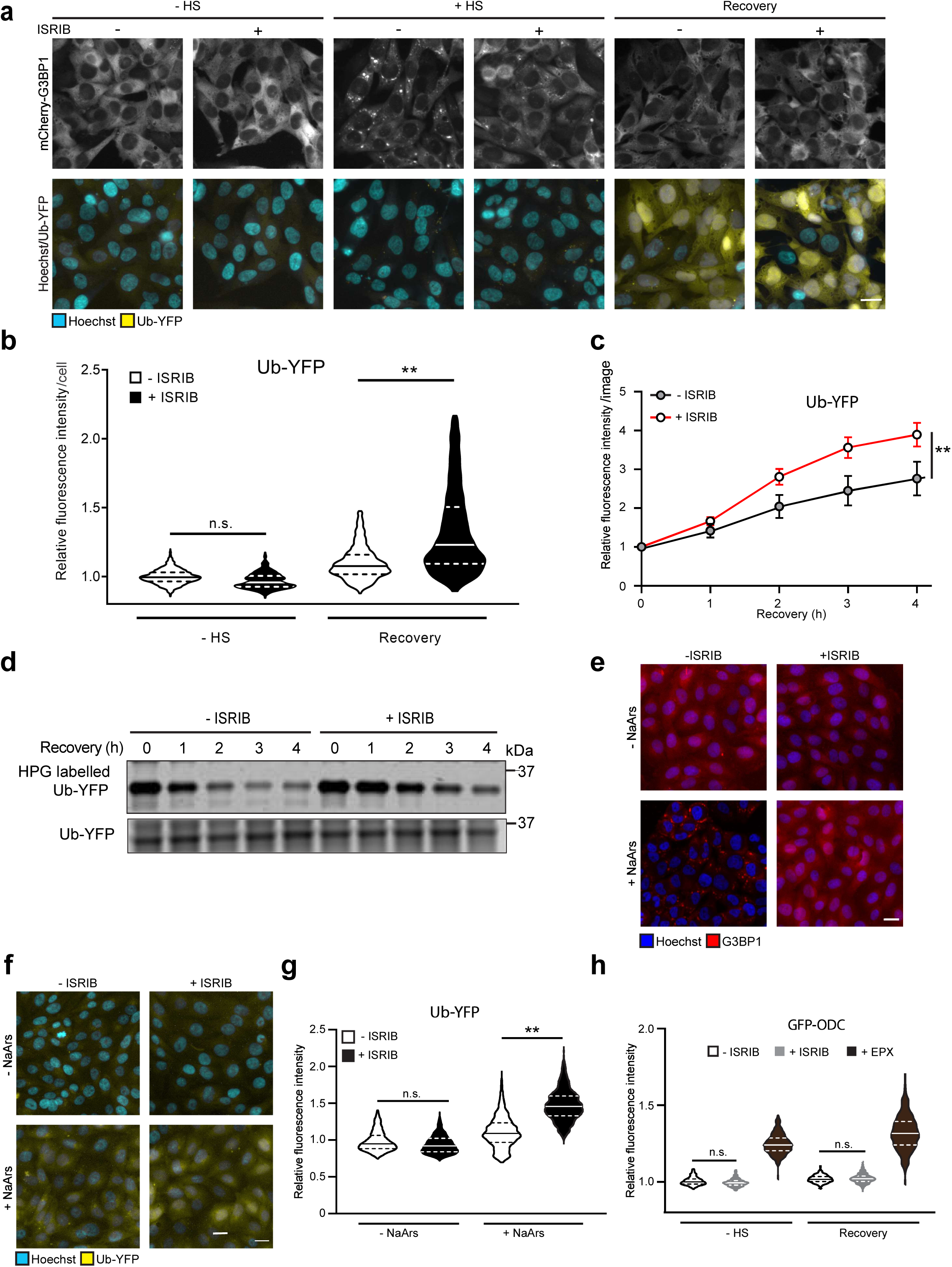
ISRIB aggravates UPS impairment during proteotoxic stress. **a)** Representative micrographs of the effect of ISRIB on Ub-YFP in thermally stressed cells. Cells were left untreated (- ISRIB)) or pretreated with ISRIB (+ ISRIB) for 30 minutes. After this initial incubation, they were left untreated (- HS) or exposed to 43°C for 30 minutes (+ HS) and followed for 4 hours after the heat shock (Recovery). Images were captured with wide-field, high-content microscope. Scale bar, 20 μm. **b)** Quantification of mean cellular YFP fluorescence intensities per cell. Experimental setup as described in (**a**). Values were normalized to untreated control cells (- ISRIB, - HS). The frequency and distribution of the relative fluorescence intensities per cell are shown as violin plots. The solid lines in each distribution represent the median, and the dash lines represent the upper and lower interquartile range limits (three independent experiments, >1000 cells analysed per condition, Kruskal-Wallis test, ** P<0.01, n.s.: not significant). **c)** Effect of ISRIB on accumulation kinetics of Ub-YFP. Kinetics of Ub-YFP accumulation in control (- ISRIB) and ISRIB-treated (+ ISRIB) MelJuSo cells in response to a 30-minutes, 43°C heat shock, followed by 4 hours at 37°C (Recovery). Data points show the mean fluorescent intensity per image with error bars indicating the standard deviation (three independent experiments, 12 images analyzed per condition, Student’s unpaired t-test, ** P<0.01). **d)** Effect of ISRIB on Ub-YFP degradation in stressed cells. MelJuSo cells expressing Ub-YFP were left untreated (- ISRIB)) or treated with ISRIB (+ ISRIB) for 30 minutes before being subjected to 43°C heat shock for 30 minutes. Pulse-chase experiment with HPG-labelled Ub-YFP. The upper panel is an in-gel detection of HPG labeled Ub- YFP in a pulse chase experiment during the recovery phase in the absence (- ISRIB) and presence of ISRIB (+ ISRIB). The total Ub-YFP reporter level was detected in the lysate by immunoblotting (lower panel). A representative result of three independent experiments is shown. **e)** Representative micrographs of MelJuSo cells expressing G3BP1-mCherry that were either pre-treated in the absence (-ISRIB) or presence (+ISRIB) of ISRIB for 30 minutes. After pretreatment, cells were treated with or without sodium arsenate (+/- NaArs) for 4 hours. Scare bar, 20 µm. **f)** Representative micrographs of the effect of ISRIB on Ub-YFP in NaArs-stressed cells. Cells were left untreated (- ISRIB) or pretreated with ISRIB (+ ISRIB) for 30 minutes. After this initial incubation they were left untreated (- NaArs) or exposed to NaArs for 4 hours (+ NaArs). Images were captured with wide-field, high-content microscope. Scale bar, 20 μm. **g)** Effect of ISRIB on Ub-YFP reporter levels in NaArs-stressed cells. Quantification of mean cellular YFP fluorescence intensities. Experimental setup as described in (D). Values were normalized to untreated control cells (- ISRIB, - HS). The frequency and distribution of the relative fluorescence intensities per cell are shown as violin plots. The solid lines in each distribution represent the median, and the dash lines represent the upper and lower interquartile range limits (three independent experiments, >1000 cells analysed per condition, Kruskal-Wallis test, ** P<0.01). **h)** Effect of ISRIB on ubiquitin-independent proteasomal degradation. Quantification of GFP intensities of MelJuSo cells expressing GFP-ODC. Cells were left untreated (- ISRIB) or incubated with ISRIB (+ ISRIB) for 30 min before being left untreated (- HS) or exposed to a 43°C heat shock for 30 minutes and followed by recovery (Recovery) for 4 hours.

Pulse-chase experiments confirmed a delayed clearance of the Ub-YFP reporter in ISRIB-treated cells that recovered from thermal stress, supporting the model that the elevated steady-state levels are a consequence of slower degradation of Ub-YFP during the recovery phase (**Fig. 1d**). The strongest effect on the turnover of Ub-YFP was observed at the first time point after the heat shock and precedes the time point at which maximal accumulation of the reporter occurred. This suggests that degradation is mostly affected during or shortly after the stress response followed by a short delay during which the reporter levels gradually start to build up.

To assess whether the same holds true for another proteotoxic condition that induce the ISR, we analyzed the effect of ISRIB on the functionality of the UPS upon induction of oxidative stress in cells exposed sodium arsenate. Treatment for 4 hours to 50 µM sodium arsenate induced the formation of stress granules in the Ub-YFP reporter cells, which was efficiently blocked by co-administration of ISRIB (**Fig. 1e**). As observed following thermal stress, the administration of ISRIB caused a larger increase in the steady-state levels of the Ub-YFP in response to sodium arsenate (**Fig. 1f,g**), suggesting that the accumulation of UPS substrates is a general consequence of ISR inhibition in stressed cells.

Impairment of the UPS in cells recovering from thermal stress is confined to ubiquitin-dependent substrates^22^. To test if the same applies to cells exposed to ISRIB, we analyzed the steady-state levels of a green fluorescent protein (GFP) reporter containing the degradation signal of ornithine decarboxylase (ODC), a ubiquitin-independent substrate of the UPS^26^. The GFP-ODC levels were not increased in thermally stressed cells regardless of the absence or presence of ISRIB, whereas inhibition of the proteasome by epoxomicin caused accumulation of this substrate (**Fig. 1h, Fig. S3**). This shows that the effect of ISRIB is confined to the degradation of ubiquitylated substrates and suggests that there is sufficient proteasome activity in ISRIB-treated cells to execute degradation. Consistent with the preserved proteasome function, ISRIB did not have a significant effect on the enzymatic activity of proteasomes in thermally stressed cells (**Fig. S4**).

### ISRIB causes accumulation and sequestration of proteasome substrates in the cytosolic, but not nuclear, compartment

We have shown previously that genetic inhibition of stress granule formation primarily affects the functionality of the UPS in the nuclear compartment with little effect on the UPS in the cytoplasm^27^. As ISRIB also prevents the formation of stress granules, we wondered if chemical interference with the formation of stress had a similar effect on the nuclear UPS. To address this question, we analyzed the UPS in the nuclear and cytosolic compartment using two compartment-specific proteasome substrates: NLS-GFP-CL1 and NES-GFP-CL1^28^. The 16 amino acid-long CL1 extension functions as a aggregation-prone degradation signal^29^, resulting in accumulation of the reporter in inclusions if not efficiently cleared^25,30^. Surprisingly, ISRIB pretreatment did not affect the degradation of the nuclear reporter in thermally stressed cells (**Fig. 2a,b**), but inhibited instead the degradation of the cytosolic reporter (**Fig. 2c,d**). The stabilization of the cytosolic aggregation-prone reporter was accompanied by an increase in the number of cells displaying accumulation of the reporter in the perinuclear region (**Fig. 2c,e**), a typical localization for sequestration of cytosolic protein aggregates^31,32^. Thus, in contrast to the nuclear UPS dysfunction in stress granule-deficient cells, the combined action of ISRIB on stress granule formation and protein synthesis results in impairment of ubiquitin-dependent proteasomal degradation in the cytosolic compartment.

**Figure 2.**
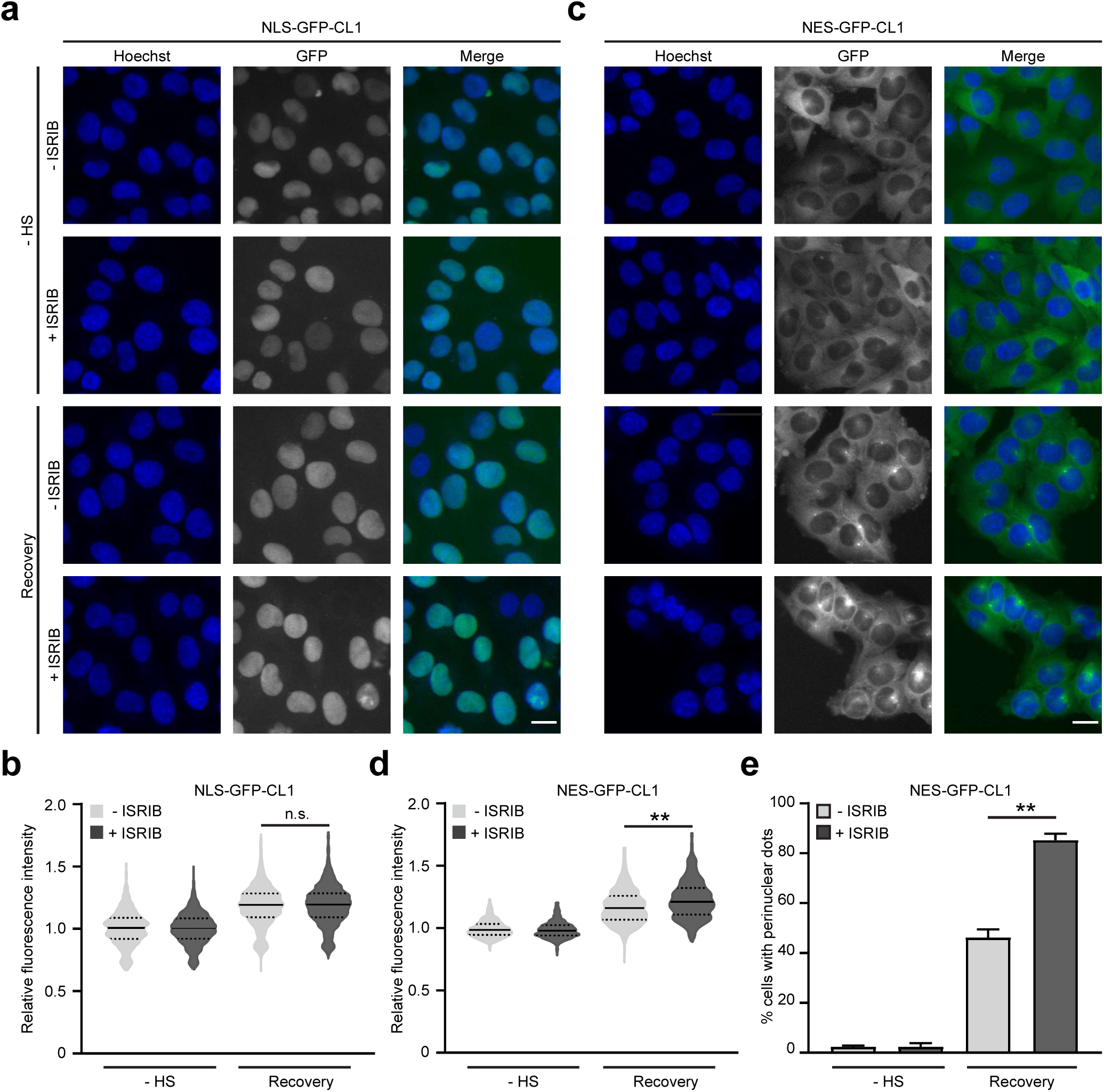
ISRIB causes accumulation and sequestration of proteasome substrates in the cytosolic, but not nuclear, compartment. **a)** Effect of ISRIB on nuclear UPS. Representative fluorescence images of MelJuSo cells expressing NLS-GFP-CL1. Cells were left untreated (- ISRIB) or incubated with ISRIB (+ ISRIB) for 30 minutes before being left untreated (- HS) or exposed to a 43°C heat shock for 30 minutes and followed by recovery (Recovery) for 4 hours. Images were captured with wide-field, high-content microscope. Scale bar, 20 μm. **b)** Quantification of mean cellular NLS-GFP-CL1 fluorescence intensities from experiment shown in (**a**). Values were normalized to untreated control cells (- ISRIB, - HS). The frequency and distribution of the relative fluorescence intensities per cell are shown as violin plots (three independent experiments, >1000 cells analysed per condition, Kruskal-Wallis test, n.s.: not significant). **c)** Effect of ISRIB on cytosolic UPS. Representative fluorescence images of MelJuSo cells expressing NES-GFP-CL1. Cells were left untreated (- ISRIB) or incubated with ISRIB (+ ISRIB) for 30 min before being left untreated (- HS) or exposed to a 43°C heat shock for 30 minutes and followed by recovery (Recovery) for 4 hours. Images were captured with wide-field, high-content microscope. Scale bar, 20 μm. **d)** Quantification of mean cellular NES-GFP-CL1 fluorescence intensities from experiment shown in (**c**). Values were normalized to untreated control cells (- ISRIB, - HS). The frequency and distribution of the relative fluorescence intensities per cell are shown as violin plots. (three independent experiments, >1000 cells analysed per condition, Kruskal-Wallis test, ** P<0.01). **e)** Percentage of cells that show cytosolic NES-GFP-CL1 foci from experiment shown in (**b**). Data represent the mean ± SD. (three independent experiments, Kruskal-Wallis test, ** P<0.01).

### ISRIB impairs the degradation of defective newly synthesized proteins (DRiPs)

As ISRIB prevents the stress-induced downregulation of protein synthesis^6^, we reasoned that treatment with this compound under proteotoxic stress conditions may increase the levels of defective newly synthesized proteins as proper co-translational folding of nascent polypeptides may become problematic. To address this question, we labelled the pool of newly synthesized peptides produced during thermal stress with puromycin. As incorporation of puromycin will terminate protein translation, this will give rise to truncated polypeptides that are commonly referred to as defective ribosome products (DRiPs), which are targeted for proteasomal degradation^33^. As anticipated based on the biological activity of ISRIB, the decrease in protein synthesis induced by thermal stress was partially reversed by ISRIB treatment (**Fig. 3a**). In untreated stressed cells, DRiPs were enriched at TIA1-positive stress granules in the cytosol and displayed, in addition, a nuclear localization (**Fig 3b, upper panels**), as reported previously^27,34^. In ISRIB-treated cells, nuclei were largely devoid of DRiPs and instead a prominent accumulation of DRiPs was observed throughout the cytosol (**Fig 3b, lower panels**).

**Figure 3.**
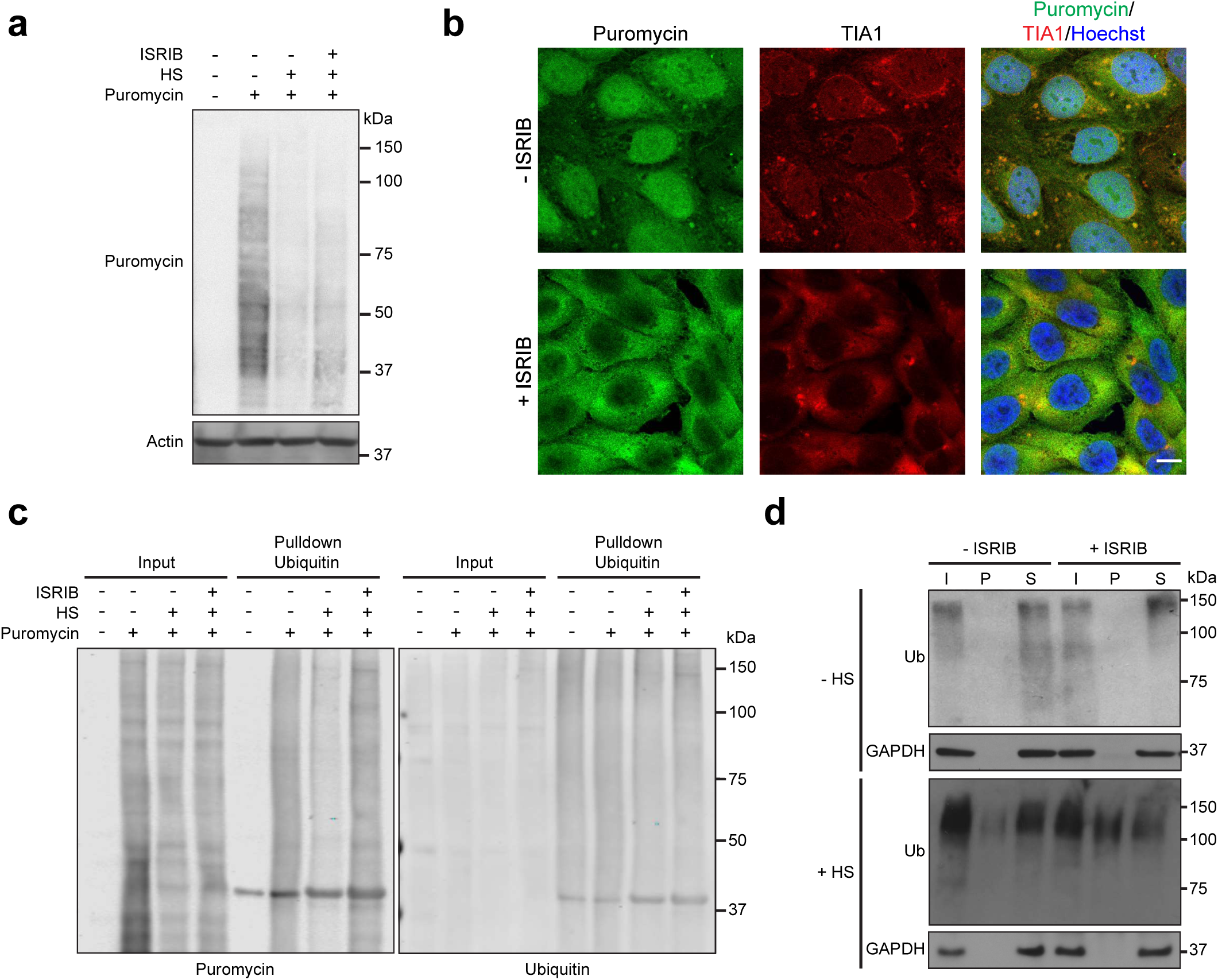
ISRIB impairs the degradation of defective newly synthesized proteins (DRiPs) **a)** Effect of ISRIB on DRiP synthesis. Analysis of puromycin incorporation in MelJuSo cells pretreated with (+ ISRIB) or without ISRIB (- ISRIB). Cells were left untreated (- HS) or exposed to 43°C for 30 minutes (+ HS). **b)** Effect of ISRIB on localization of DRiPs. Representative fluorescence images of MelJuSo cells that were pre-treated with (+ ISRIB) or without ISRIB (- ISRIB). Cells were exposed to 43°C for 30 minutes (+ HS) in the presence of puromycin. Puromycin and the stress granule marker TIA1 were visualized by immunostaining. Images were captured with laser scanning confocal microscope. Scare bar, 10 μm. **c)** Western blot analysis of ubiquitylated DRiPs in MelJuSo cells expressing Ub-YFP that were incubated in the absence of presence of ISRIB for 30 minutes before left untreated or subjected to 43°C heat shock for 30 minutes with or without 5 ug/ml puromycin. The cells were lysed as indicated and ubiquitylated proteins were pulled down with TUBEs. The input and pulldown samples were analyzed by immunoblotting with antibodies directed against ubiquitin and puromycin. **d)** Western blot analysis of Triton X-100 solubility assay of MelJuSo cells incubated with or without ISRIB for 30 minutes before left untreated (-HS) or subjected to 43°C heat shock for 30 minutes (+HS). Ubiquitylated proteins were detected in total lysates (I), insoluble pellet fraction (P), and soluble fraction (S). GAPDH is used as a control for soluble proteins.

To investigate the levels of ubiquitylated DRiPs, Tandem Ubiquitin Binding Entities (TUBEs) were used to pulldown polyubiquitylated, puromycin-labelled proteins from the lysate of cells exposed to heat shock in the absence or presence of ISRIB. Indeed, ISRIB treatment promoted the accumulation of ubiquitylated DRiPs in stressed cells (**Fig. 3c**). In line with the assumption that the persistent translation under stress conditions increased the load of misfolded or unfolded proteasome substrates, the administration of ISRIB to cells exposed to heat shock increased the pool of Triton X-100 insoluble ubiquitylated proteins (**Fig. 3d**). These results support a model where persistent protein synthesis during proteotoxic stress causes cytosolic accumulation of ubiquitylated proteins that are prone to aggregation.

### Ribosome quality control (RQC) substrates overload the UPS in ISRIB-treated cells

As DRiPs are typically targeted for proteasomal degradation by the ribosome quality control (RQC) machinery^35^, we wondered if the increase in DRiPs in ISRIB-treated cells had an impact on the ability of cells to efficiently clear RQC substrates. We first checked if RQC is affected by ISRIB by analyzing the effect of ISRIB-treatment on ribosome stalling at poly(A) sequences using a poly(A) read-through reporter^36^. This showed that ISRIB treatment did not interfere with the initial steps of the RQC leading to persistent ribosome stalling and disassembly since the ribosomes failed to read through the poly(A) sequence (**Fig. 4a,b**). To analyze the fate of polypeptides intercepted by RQC, we used a GFP^nonstop^ reporter, that due to a mutation of the GFP stop codon, produces a GFP-tagged readthrough RQC substrate that can be readily quantified^37^. We found that ISRIB caused an increase in GFP^nonstop^ levels that were comparable with the levels reached upon treatment with the proteasome inhibitor epoxomicin (**Fig. 4c,d**). Co-treatment of ISRIB and epoxomicin did not cause a further accumulation of GFP^nonstop^, indicating that the accumulation of GFP^nonstop^ in stressed, ISRIB-treated cells was not due to an increase in synthesis of the reporter but caused by a delayed clearance of GFP^nonstop^ by the UPS. This also indicates that the effect of ISRIB on degradation of RQC substrates is stronger than the soluble Ub-YFP reporter. Thus, the increase in defective newly synthesized proteins is accompanied by inefficient clearance of RQC substrates by the UPS.

**Figure 4.**
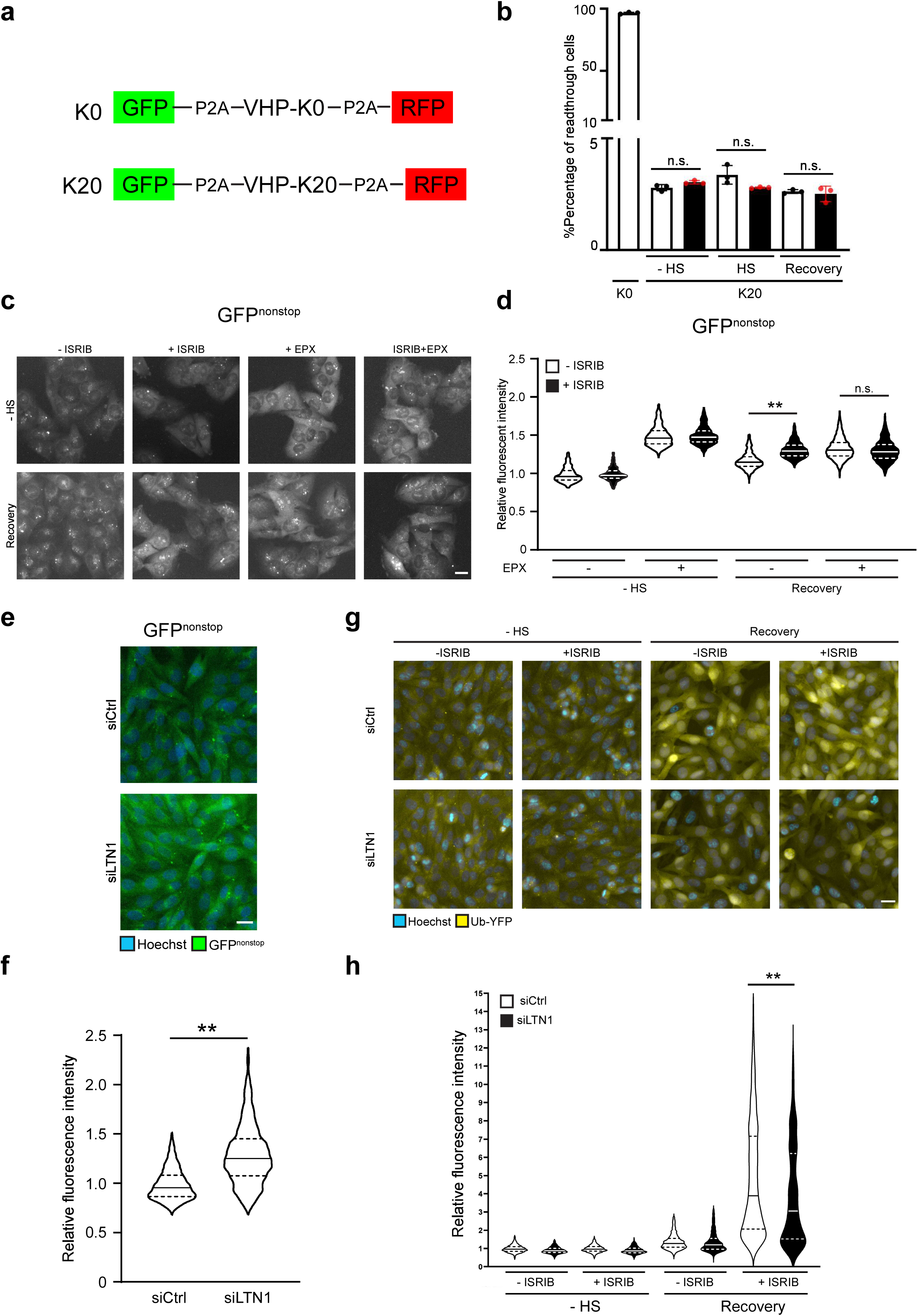
Ribosome quality control (RQC) substrates overload the UPS in ISRIB-treated cells. **a)** Schematic representation of the readthrough double reporters GFP-P2A-K0-P2ARFP-(K0) and GFP-P2A-K20-P2A-RFP (K20). **b)** Flow cytometric quantification of MelJuSo cells transfected with pmGFP-P2A-K20-P2A-RFP (K0) or pmGFP-P2A-K20-P2A-RFP (K20) respectively. Cells expressing GFP-K20-RFP were either pre-treated in the absence (-ISRIB) or presence (+ ISRIB) of ISRIB for 30 minutes. After pretreatment, cells were left untreated (-HS), subjected to 43°C heat shock (+HS) or followed by 4 hours recovery (Recovery). Readthrough region was gated in K0 transfected cells, exhibiting linear correlation between GFP and RFP fluorescence. Data represent the mean ± SD. (three independent experiments, n.s.: not significant). **c)** Effect of UPS on RQC substrate GFP^nonstop^. Representative fluorescence images of MelJuSo cells expressing GFP^nonstop^. Cells were left untreated (- ISRIB), incubated with ISRIB for 30 minutes (+ ISRIB), incubated with proteasome inhibitor epoxomicin for 4 hours (+ EPX) or combined ISRIB with proteasome inhibitor epoxomicin (ISRIB+EPX) before being left untreated (- HS) or exposed to a 43°C heat shock for 30 minutes and followed by recovery (Recovery) for 4 hours. Images were captured with wide-field, high-content microscope. Scale bar, 20 μm. **d)** Quantification of mean cellular YFP fluorescence intensities in MelJuSo cells stably expressing GFP^nonstop^. Cells were pre-treated with or without ISRIB for 30 minutes and either left untreated (- HS) or 43°C heat shock for 30 minutes and followed by 4 hours (Recovery), in presence or absence of the proteasome inhibitor epoxomicin (+ EPX). Values were normalized to untreated control cells (- ISRIB, - HS). The frequency and distribution of the relative fluorescence intensities per cell are shown as violin plots. (three independent experiments, >1000 cells analysed per condition, Kruskal-Wallis test, ** P<0.01, n.s.: not significant). **e)** Listerin depletion impairs RQC. Representative micrographs of MelJuSo cells expressing GFP^nonstop^ that were either transfected with control siRNA (siCtrl) or LTN1 siRNA (siLTN1). Images were captured with wide-field, high-content microscope. Scare bar, 20 μm. **f)** Quantification of mean cellular GFP fluorescence intensities from experiment shown in (A). Values were normalized to transfection control cells (siCtrl). The frequency and distribution of the relative fluorescence intensities per cell are shown as violin plots. (three independent experiments, >1000 cells analysed per condition, Kruskal-Wallis test,** P<0.01). **g)** Effect of listerin depletion on ISRIB effect on Ub-YFP. Representative micrographs of MelJuSo cells expressing Ub-YFP that were either transfected with control siRNA (siCtrl) or LTN1 siRNA (siLTN1). After transfection, cells were left untreated (-ISRIB) or pretreated with ISRIB (+ISRIB) for 30 minutes. After this initial incubation they were left untreated (- HS) or exposed to 43°C for 30 minutes and followed for 4 hours after the heat shock (Recovery). Images were captured with wide-field, high-content microscope. Scale bar, 20 μm. **h)** Quantification of mean cellular YFP fluorescence intensities. Experimental setup as described in (C). Values were normalized to untreated control cells (- ISRIB, - HS). The frequency and distribution of the relative fluorescence intensities per cell are shown as violin plots. The solid lines in each distribution represent the median, and the dash lines represent the upper and lower interquartile range limits (three independent experiments, >700 cells analysed per condition, Kruskal-Wallis test, *** P=0.0007). One representative experiment out of three.

We hypothesized that the impaired degradation of reporter substrates was a consequence of the increased levels of RQC substrates in ISRIB-treated cells, which overwhelm the cytosolic UPS and compete with other ubiquitylated substrates for degradation. The ubiquitin ligase listerin is part of the RQC complex and targets aberrant nascent chains for ubiquitin-dependent proteasomal degradation ^38^. To test our hypothesis, we analyzed the effect of siRNA depletion of listerin on the UPS activity of stressed cells in the absence or presence of ISRIB. Western blotting confirmed that listerin levels were reduced by siRNA (**Fig. S5**) and, importantly, that this resulted in impaired RQC as evidenced by a significant increase in the GFP^nonstop^ reporter (**Fig. 4e,f**). Interestingly, we observed that listerin depletion significantly reduced the accumulation of Ub-YFP in thermally stressed ISRIB-treated cells (**Fig. 4g,h**). These data are consistent with a model where the RQC is an important source of ubiquitylated substrates that overload the UPS in ISRIB-treated, stressed cells. Thus, these data suggest that the persistence of protein synthesis in ISRIB-treated cells exposed to proteotoxic stress leads to the *de novo* production of defective proteins that are intercepted by the RQC and accumulate in the cytosol, thereby impairing the functionality of the UPS.

## Discussion

In summary, we show that ISRIB compromises ubiquitin-dependent proteasomal degradation in cells that undergo proteotoxic stress. Our data point to the persistent protein synthesis during proteotoxic stress as the primary cause of this effect as ISRIB blunts the protective inhibition of protein synthesis, which is normally triggered by the ISR^39^. The reduction of protein synthesis during proteotoxic stress is best understood as an attempt to minimize the accumulation of aggregation-prone misfolded proteins, as nascent chains will be hindered from adopting their native conformation under challenging conditions^39^. Chemical inhibition of the ISR is likely to increase the load of newly synthesized proteins that need to be intercepted by the RQC and thereby targeted for proteasomal degradation. The partial increase in the levels of newly synthesized proteins that we observed in ISRIB-treated stressed cells is likely to be an underestimation as a large pool of these proteins may be co-translationally degraded, thereby being a potential source for the overload of the RQC pathway. The interplay between the ISR and RQC may even further enhance UPS overload as the inability to activate the ISR has been shown to boost protein ubiquitylation by RQC in yeast^40^. While our data suggest that saturation of the UPS with RQC substrates plays an important role, we cannot exclude the possibility that ISRIB treatment also compromises the UPS at other levels, such as modulating expression of regulators of this proteolytic system.

Our data indicate that in ISRIB-treated cells degradation of ubiquitin-dependent, but not ubiquitin-independent, substrates is compromised. We have previously shown that impairment of the UPS during proteotoxic stress is not caused by saturation of the capacity of proteasomal activity in cells^22^. Instead, the available pool of free ubiquitin appears to a limiting factor, which gives rise to competition between substrates for ubiquitin tagging, a demand that will dramatically increase during proteotoxic stress due to the accumulation of misfolded proteins^22^. Our finding that that activity of the proteasome is not reduced in ISRIB-treated cells, which is also supported by the unabated degradation of a ubiquitin-independent substrate, is consistent with an increased demand on ubiquitylation as the primary cause for the accumulation of UPS substrates. We have speculated that the competition for ubiquitin may be important to provide crosstalk between the various proteolytic and non-proteolytic processes that critically depend on ubiquitin conjugation^41,42^. It is therefore possible that ISRIB will also indirectly compromise other ubiquitin-dependent processes, such as transcription and DNA repair, which may have a further negative impact on the viability of cells. As the potential negative consequences on these cellular processes may become a source for adverse effects, it will be important to further explore the effects of ISRIB on other ubiquitin-dependent processes.

Interestingly, the inhibitory effect of ISRIB on the degradation of an RQC reporter substrate was the strongest with the effect on the degradation of soluble reporter substrates being less severe. This suggests that the competition is fiercest at the level of ubiquitylation of RQC substrates and may not only involve competition for free ubiquitin but also competition for various enzymes involved in ubiquitylation of RQC substrates. The effect on soluble substrates may be caused by depletion of free ubiquitin and/or flooding of the UPS with ubiquitylated RQC substrates. The general effect on ubiquitin-dependent proteasomal degradation may also explain the original observation that ISRIB treatment compromises the ability of cells to cope with endoplasmic reticulum (ER) stress^8^, as dealing with this condition requires efficient degradation of misfolded ER proteins through ubiquitin-dependent ER-associated degradation (ERAD) by the UPS^43^. Notably, ER stress itself has already been found to push the UPS to its limits^25^, and further havoc of the UPS may be the reason for the impaired ability of ISRIB-treated cells to adapt to ER stress.

The ISR appears to be a double-edged sword as its intrinsic protective effect on the detrimental consequence of proteotoxic stress goes hand in hand with its contribution to the etiology of a number of diseases that are associated with aberrant activation of this stress reponse^1^. In support of an overall beneficial effect of ISRIB, it has been shown that ISRIB can readily cross the blood-brain barrier with good pharmacokinetic properties and enhance long-term memory in mice^8^. Moreover, the capacity of ISRIB to prevent the formation of stress granules may have additional therapeutic potential since these structures have been linked to several neurodegenerative diseases, most notably amyotrophic lateral sclerosis^10^. On the other hand, prolonging the ISR by pharmacologically inhibition of the dephosphorylation of eIF2α, which results in a delay in translational recovery, prevents physiological and molecular defects in mouse models for Charcot-Marie-Tooth 1B and amyotrophic lateral sclerosis, suggesting a general beneficial effect in protein-misfolding diseases^16,17^. Maintaining efficient protein quality control by preserving UPS activity through extending the time window for ISR-induced translational inhibition may contribute to the protective effect of these compounds.

Whether the effect of ISRIB and other ISR inhibitors will be beneficial or detrimental may depend on the ability of such compounds to prevent the aberrant activation of the ISR without hindering its protective function. Earlier studies suggest that ISRIB may meet this requirement as the robust activation of the ISR in response to severe stress, is not compromised by ISRIB whereas the lingering aberrant response associated with pathologies is efficiently blunted^6,44^. Our data suggest that, even though the use of ISRIB may be promising in therapeutic settings, caution should be practiced as its negative impact on the clearance of misfolded proteins by the UPS may exacerbate conditions that favor protein aggregation. In particular, chronic curtailing of the ISR in neurons can turn out to be problematic since the accompanying UPS impairment may accelerate the age-dependent accumulation of aggregation-prone proteins that are linked to neurodegeneration^45^.

## Material and Methods

### Plasmids

The GFP^nonstop^ expression plasmid was generated by PCR amplifying the GFP open reading frame from EGFP-C1 using primers 5’-CGA TCG ACT AGT ACG CGT GTT ACA AAT AAA GCA ATA GCA TCA-3’ (forward primer) and 5’-ACG CGT ACT AGT CGA TCG CTT GTA CAG CTC GTC CAT G-3’ (reverse primer) and using NEBuilder HiFi DNA Assembly Master Mix (New England Biolabs) according to the manufacturer’s instructions. The NLS-GFP-NLS-GFP-CL1 and NES-GFP-NES-GFP-CL1 ^28^, was a gift from Dr. Ron Kopito (Stanford University). The pmGFP-P2A-K0-P2A-RFP (Addgene #105686) and pmGFP-P2A-K20-P2A-RFP (Addgene #105688) have been described previously ^36^.

### Cell culture and transfection

The human melanoma MelJuSo cell lines (RRID:CVCL_1403) were cultured in DMEM+GlutaMAX (Life Technologies) supplemented with 10% Fetal Bovine Serum (FBS) in a at 37 °C and 5% CO_2_. Cell lines are routinely tested for mycoplasma infection. Generation of MelJuSo Ub-YFP stable cell lines have been described previously ^25^. The Mel JuSo GFP-ODC, NLS-GFP-NLS-GFP-CL1, NES-GFP-NES-GFP-CL1 and GFP^nonstop^ cell lines were created by transfection with corresponding GFP-ODC, NLS-GFP-NLS-GFP-CL1, NES-GFP-NES-GFP-CL1 and GFP^nonstop^ plasmids using Lipofectamine 3000 (Life Technologies) respectively. Clones were selected in the presence of 1.5 mg/ml G418 (Gibco). A bacterial artificial chromosome (BAC) containing the genomic sequence of the stress granule marker G3BP1, N-terminally tagged with mCherry via homologous recombination, was stably transfected into the MelJuSo Ub-YFP cell line ^46^.Transient transfection was performed using Lipofectamine 3000 (Life Technologies) according to the manufacturer’s instructions. For knockdown experiments, the following siRNA constructs were purchased from Life Technologies: control siRNA (4390843), LTN1 siRNA (S25003). Lipofectamine RNAiMax (Life Technologies) was used for transfection of siRNA according to the manufacturer’s instructions.

### Chemical treatment

Before subjected to heat shock, the ISRIB treatment group cells were pre-treated with 200 nM ISRIB (Merk) for 30 minutes. After heat shock, 100 nM epoxomicin (Sigma) was added for 4 hours. To test the effect of ISRIB on sodium arsenate-induced stress, cells were first pretreated with 200 nM ISRIB for 30 min followed by incubation with 50 µM sodium arsenate (Sigma) for 4 hours. For the proteasome inhibitor titration experiments, Ub-YFP MelJuSo cells were treated with the indicated epoxomicin concentration for 4 hours.

### Immunofluorescence

MelJuSo cell lines were grown on cover slips overnight and treated as indicated. Cells were fixed using 4% paraformaldehyde (Life Technologies) for 15 min, permeabilized using 0.2% Triton-X100 in 1x phosphate buffered saline (PBS) for 15 min and blocked using 3% BSA (Sigma) in 1x PBS for 30 min. The primary antibodies were diluted in 0.1% Tween-20 in PBS and incubated overnight at 4°C. The following primary antibodies were used: anti-puromycin (MABE343; Sigma), anti-TIA1 (#140595; Abcam). Goat anti-rabbit IgG or goat anti-mouse IgG coupled to AlexaFluor 488 or 568 (Life Technologies) were diluted 1:500 in 0.1% tween 20 in PBS. Nuclear staining was performed using Hoechst 33342 (Molecular Probes) 1:5000 in PBS for 15 min. Fixed cells were examined with a Zeiss LSM 880 confocal laser scanning microscope (Plan-Neofluar 63x/1.4 oil objective). Image processing was performed with FiJi and quantitative analyses were performed using cell profile software.

### Immunoblotting

Equal amounts of cells were lysed in 1X LDS sample buffer (Life Technologies) containing 10% NuPAGE reducing agent (Life Technologies) and lysates were boiled at 95°C for 5 minutes. Cell protein extracts were resolved by Bis-Tris polyacrylamide gel electrophoresis gels (Life Technologies) and run in 1 X MOPS buffer (Life Technologies). Proteins were transferred onto nitrocellulose membranes (GE Healthcare) in a Tris-glycine transfer buffer containing 20% methanol. After blocking in Tris-buffered saline (TBS)/non-fat milk 5% containing 0.1% Tween-20, membranes were incubated with primary antibodies. The following antibodies were used: anti-GFP (Abcam, 290), anti-GAPDH (Abcam, 9485), anti-β-actin (Abcam, 8226), anti-puromycin (Sigma, MABE343), anti-ubiquitin (Santa Cruz, sc-8017), anti-LTN1 (Proteintech, 28452-1-AP), anti-eIF2α(pSer51) (Cell Signaling, 3389) and eIF2α (Cell Signaling, 2103). After incubation with the primary antibody, the membranes were washed with TBS-Tween-20 0.1% and incubated with secondary goat anti-rabbit or anti-mouse horseradish peroxidase (HRP)-linked antibodies. Detection was performed by enhanced chemiluminescence (Amersham ECL reagents, GE Healthcare) on Medical X-ray films (Fujifilm). Alternatively, secondary antibodies coupled to near-infrared fluorescent dyes (LI-COR) were used, and membranes scanned with an Odyssey scanner (LI-COR) and analyzed with Image Studio Lite analysis software version 5.2 (LI-COR).

### Pulse-chase analysis

Cells were seeded overnight in a 6 well plate. Thirty min before thermal stress was induced, methionine was depleted with methionine-free RPMI medium (Life Technologies) simultaneously pre-treating with ISRIB. After heat shock, the medium was replaced with methionine-free RPMI Medium containing 50 µM of the methionine analogue L-homopropargylglycine (HPG) (Jena Bioscience, CLK-1067) for incorporation of HPG during protein synthesis during the recovery phase. After washing cells once in PBS, lysis was performed for 30 min using 250 µl/sample. After 10 minutes centrifugation at 4°C and 16,000g, supernatant was used to perform Click chemistry for 30 minutes using 10 µM 800CW Azide Infrared Dye (LI-COR, CA1007-02). Dilution buffer was added to dilute the detergent concentration before the immunoprecipitation (IP) was performed with equilibrated GFP-Trap A beads by end-over-end tumble for 1.5 hours at 4°C. Collection of beads bound material and following washing steps with lysis or dilution buffer were done by centrifugation at 4°C and 2500g for 2 minutes discarding the supernatant. Next, 2x SDS sample buffer with reducing agent was added to beads before boiling samples for 10 minutes at 95°C to dissociate immune complexes. Prior proceeding with SDS-PAGE and western blotting, beads were spun down at 2500g for 1 minute. Supernatant was loaded on the gel. In-gel detection of fluorescent labelled HPG was performed with an 800 nm laser was performed with an Odyssey scanner (LI-COR). The gel was next transferred to a nitrocellulose membrane for detection of total Ub-YFP levels by western blot analysis.

### TUBE pulldown

After indicated treatment, cells were harvested and lysed in RIPA buffer on ice for 30 minutes. After 10 minutes centrifugation at 4°C and 16000g, supernatant was mixed with pre-equilibrium TUBE beads (Life Sensors) and incubated overnight at 4°C. The beads were washed three times with detergent free RIPA buffer. After the last wash, 2x SDS sample buffer with reducing agent was added to beads before boiling samples for 10 minutes at 95°C to dissociate immune complexes. Prior proceeding with SDS-PAGE and western blotting, beads were spun down at 2500g for 1 min. Supernatant was loaded on the gel.

### High-content microscopy

For live imaging, MelJuSo Ub-YFP were seeded in 96 wells plate (Falcon) at 4,000 cells per well and incubated overnight. After indicated treatment, the medium was replaced by Leibovitz’s L-15 medium (Life Technologies). Four sites per well were imaged with 10 minutes intervals using an automated widefield microscope (ImageXpress Micro; Molecular Devices) at 37 °C. The fluorescent intensity was quantified non-blinded with the CellProfiler software in which nuclear and cytoplasmic segmentations were determined by intensity thresholding of the Hoechst and GFP signals. Fluorescence intensities of YFP and GFP were measured in the nuclear segmentations for all experiments, except for NES-GFP-CL, for which the cytoplasmic intensities were measured.

### Proteasome activity

MelJuSo expressing Ub-YFP cells were treated as indicated and harvested in lysis buffer (25 mM HEPES pH 7.2, 50 mM NaCl, 1 mM MgCl_2_, 1 mM ATP, 1 mM DTT, 10% glycerol, 1% Triton X-100). After centrifugation, protein concentrations were measured with the protein assay dye reagent (Bio-Rad) and 10 μg protein was mixed with 80 μl reaction buffer (lysis buffer without Triton X-100) and 10 μl suc-LLVY-AMC (Affiniti, P802) for a final concentration of 1 μM suc-LLVY-AMC. As a control, 100 nM of the proteasome inhibitor epoxomicin (Sigma) was added to the reaction mixture. Samples were analyzed in a microplate reader (FLUOStar OPTIMA) at 355 nm/460 nm every minute for 1 hour.

### Triton X-100 solubility assay

MelJuSo expressing Ub-YFP cells were seeded in 6 well plates and incubated overnight. Prior to lysis, cells were either pretreated with DMSO or ISRIB for 30 minutes and followed by either left untreated or exposed to 43°C for 30 min. Cells were washed twice in cold PBS, followed by scraping in 400 µl lysis buffer on ice (1% Triton X-100 (Sigma), 1x complete EDTA-free Protease Inhibitor Cocktail (Roche), 20 mM N-Ethylmaleimide (Sigma), in PBS). Lysates were left on ice for 30 minutes, 200 uL of lysates were added with SDS-PAGE loading buffer and followed by 5 minutes of boiling at 95°C (Input fraction). Two hundred µl of lysates were followed by centrifugation (12,000 g, 4°C, 10 minutes). SDS-PAGE loading buffer was added to the supernatants, followed by 5 minutes of boiling at 95°C (Triton X-100 soluble fraction). The pellet was resuspended in loading buffer and followed by boiling for 10 min at 95°C (Triton X-100 insoluble fraction).

### Translation readthrough assay

MelJuSo cells were seeded in multi wells plates and incubated overnight. The cells were transfected with the pmGFP-P2A-K0-P2A-RFP (K0) or pmGFP-P2A-K20-P2A-RFP (K20) for 24 hours. The transfected cells were treated with or without ISRIB for 30 minutes and either left without treatment (-HS), heat shock for 30 minutes (+HS) or heat shock for 30 minutes and followed by 4 hours recovery (Recovery). The GFP and RFP fluorescent intensities were detected by flow cytometry (BD LSR Ⅱ) and data was analyzed with the FlowJo.

### Statistical analysis

Statistical analyses were performed using GraphPad Prism version 8.3. To test for Gaussian distribution, the D’Agostino & Pearson or Shapiro-Wilk normality test (for smaller sample sizes) were used. If the normality test was passed, data were analyzed by Student’s unpaired t-test (two groups) or by ANOVA (more than two groups). If the data were not normally distributed, statistical analysis was performed using the nonparametric Mann-Whitney test (two groups) or Kruskal-Wallis test for multiple comparisons, with Dunnett’s or Tukey’s test to adjust for multiple comparisons. Grubbs’ test was used for the detection of outliers. Adjusted p-values are shown. Data are shown as mean ± SD (standard deviation), unless stated otherwise, as indicated in each figure legend. The following p-values were considered significant: * P≤0.05; ** P≤0.01.

## Acknowledgements

We thank Vanessa Aires-Mofreita for practical help, Maria Masucci and the members of the Dantuma lab and Masucci lab for helpful input and the Biomedicum Imaging Center (BIC) and Biomedicum Flow Cytometry Core Facility for their assistance. This work was supported by the Swedish Research Council (N.P.D. 2021-02562), the Swedish Cancer Society (N.P.D. CAN 211653Pj), the Joint Programme Neurodegenerative Diseases (JPND) (S.A., N.P.D.), the Chinese Scholarship Council (S.X.) and the Karolinska Institute (KID grant, E.B). M.E.G. was supported by research fellowships from the Deutsche Forschungsgemeinschaft (DFG) (GI-1329/1-1). The Dantuma lab is member of the COST network ProteoCure.

## Author contributions

**S.X., S.A., F.A.S., N.P.D.** conceptualized the study; **S.X., M.E.G, E.B.** performed and analyzed experiments; **I.P.** generated stable cell lines; **F.A.S.** assisted in microscopy and image analysis; **S.X., N.P.D.** wrote the manuscript; **S.A., N.P.D.** coordinated the project; all authors edited and approved the final manuscript.

## Data Availability

This study does not contain deposited data. Original data can be provided upon request.

