## Supplementary Figures for "Chemical inhibition of the integrated stress response impairs the ubiquitin-proteasome system"

### Figure S1 Induction of ISR by heat shock and

MelJuSo were pretreated in the absence (-ISRIB) or presence (+ ISRIB) of ISRIB for 30 minutes and followed by either left untreated (- HS) or exposed to 43°C for 30 minutes (HS). Cell lysates were probed for phosphorylated eIF2α (pSer51), total eIF2α and beta-actin.

### Figure S2 Accumulation of Ub-YFP in response to proteasome inhibition.

MelJuSo expressing Ub-YFP and mCherry-G3BP1 were treated for 4 hours with increasing concentrations of the proteasome inhibitor epoxomicin (EPX). For each condition, the mean fluorescence intensity of at least 7000 cells was quantified. The frequency and distribution of the relative YFP fluorescence intensities per cell are shown as violin plots. The solid lines in each distribution represent the median, and the dash lines represent the upper and lower interquartile range limits (>7000 cells analysed per condition, Kruskal-Wallis test, **P<0.0001). One representative experiment out of two.

### Figure S3 ISRIB does not affect ubiquitin-independent degradation.

Effect of ISRIB on ubiquitin-independent proteasomal degradation. Representative fluorescence images of MelJuSo cells expressing GFP-ODC. Cells were left untreated (- ISRIB)) or incubated with ISRIB (+ ISRIB) for 30 min before being left untreated (-HS) or exposed to a 43°C heat shock for 30 minutes and followed by recovery (Recovery) for 4 hours. Images were captured with wide-field, high-content microscope. Scale bar, 20 µm.

### Figure S4 Proteasome activity is preserved in ISRIB-treated cells.

MelJuSo expressing Ub-YFP cells were pretreated in the absence (-ISRIB) or presence (+ ISRIB) of ISRIB for 30 minutes and followed by either left untreated (- HS), exposed to 43°C for 30 minutes and followed for 4 hours after heat shock (Recovery).

The chymotrypsin activity (β5 subunit) of the proteasome was detected by following conversion of the fluorogenic Suc-LLVY-AMC substrate over 1 hour. As a control, 100 nM proteasome inhibitor epoxomicin (EPX) was added to the reaction mixture to inhibit proteasome activity. Data represent the mean ± SD. (n.s.: not significant).

**Figure S5 Listerin siRNA depletion.**

MelJuSo cells expressing Ub-YFP were transfected with control siRNA or siLTN1 siRNA. Cell lysates were analyzed by immunoblot with LTN1 and GAPDH antibodies.

Xu et al, 2023 Figure S1

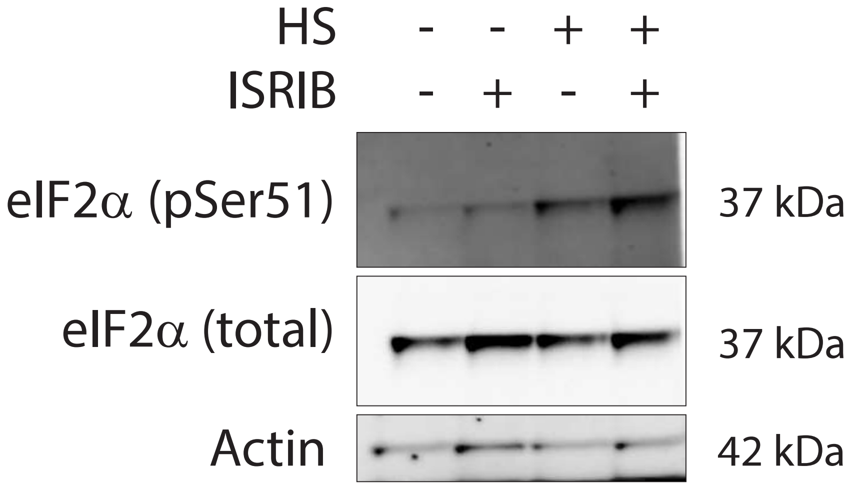

Xu et al, 2023 Fig S2

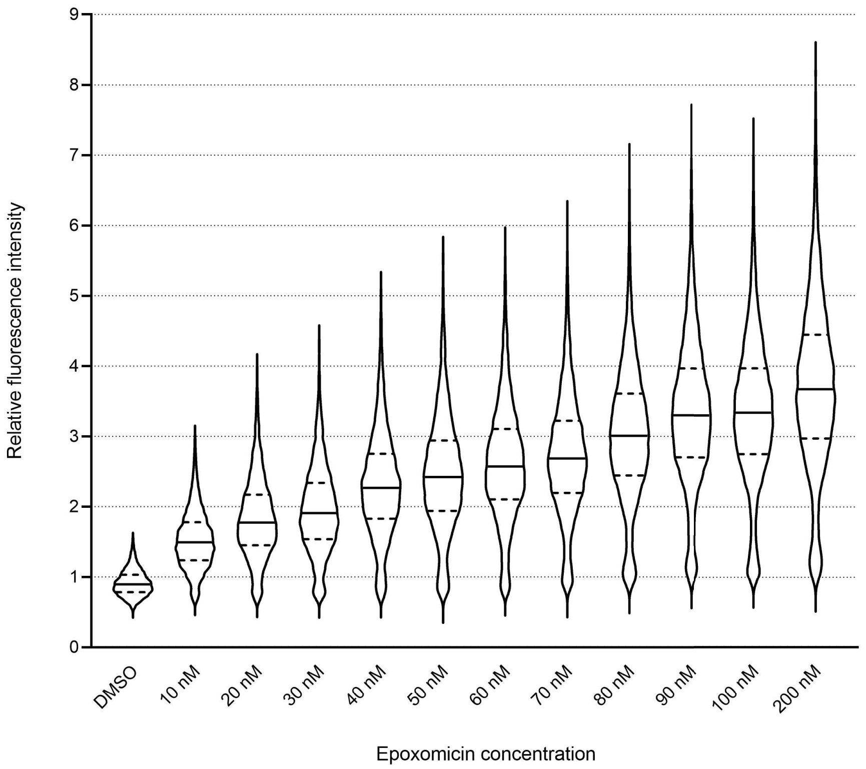

Xu et al, 2023 Figure S3

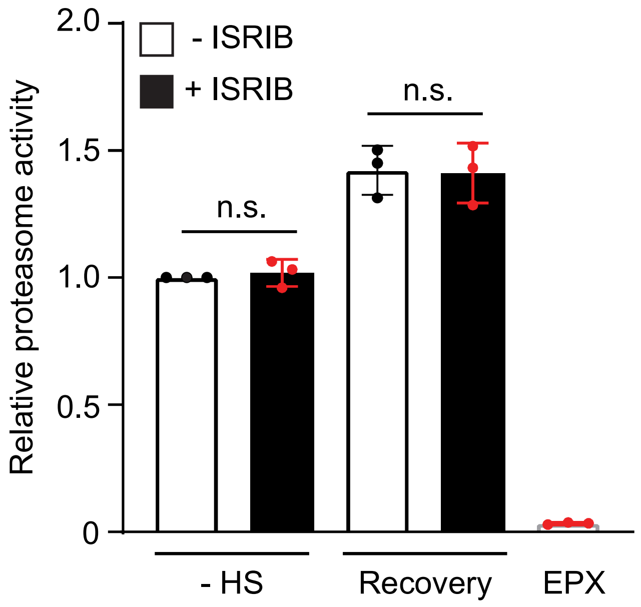

Xu et al, 2023 Figure S4

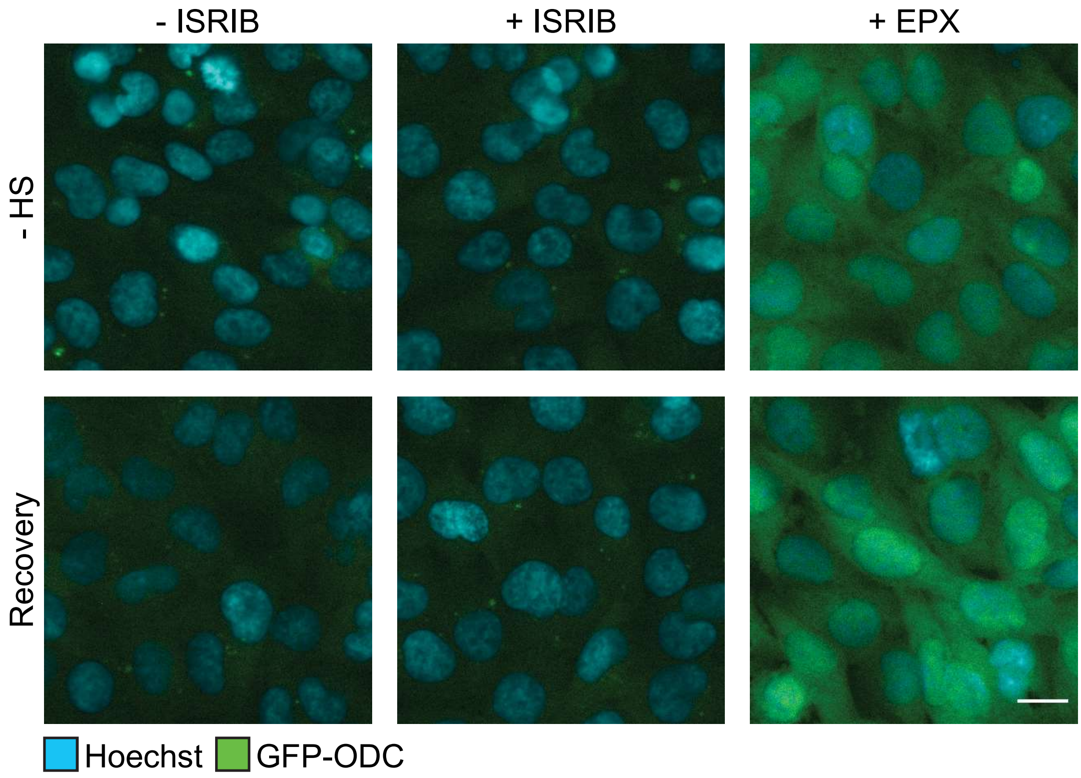

Xu et al, 2023 Figure S5

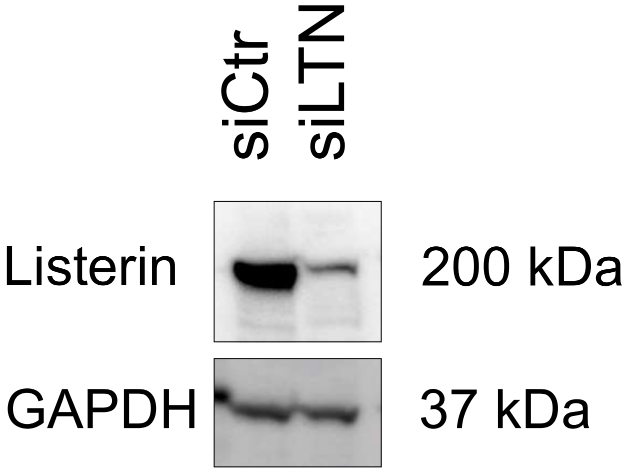
